# An entorhinal-like region in food-caching birds

**DOI:** 10.1101/2023.01.05.522940

**Authors:** Marissa C. Applegate, Konstantin S. Gutnichenko, Emily L. Mackevicius, Dmitriy Aronov

## Abstract

The mammalian entorhinal cortex routes inputs from diverse sources into the hippocampus. This information is mixed and expressed in the activity of many specialized entorhinal cell types, which are considered indispensable for hippocampal function. However, functionally similar hippocampi exist even in non-mammals that lack an obvious entorhinal cortex, or generally any layered cortex. To address this dilemma, we mapped extrinsic hippocampal connections in chickadees, whose hippocampi are used for remembering numerous food caches. We found a well-delineated structure in these birds that is topologically similar to the entorhinal cortex and interfaces between the hippocampus and other pallial regions. Recordings of this structure revealed entorhinal-like activity, including border and multi-field grid-like cells. These cells were localized to the subregion predicted by anatomical mapping to match the dorsomedial entorhinal cortex. Our findings uncover an anatomical and physiological equivalence of vastly different brains, suggesting a fundamental nature of entorhinal-like computations for hippocampal function.

## INTRODUCTION

A defining feature of episodic memories is that they bind together different types of internal and external information.^1,2^ In the mammalian brain these streams of information from different parts of the brain are not conveyed directly to the hippocampus, a site of memory storage. Instead, signals first converge in the entorhinal cortex – a part of the hippocampal formation that in turn provides input to the hippocampus itself.^3,4^ Consistent with these convergent signals, entorhinal neurons exhibit a remarkable diversity of firing patterns that encode behaviorally-relevant variables like location, speed, head direction, environmental boundaries, sensory cues, objects, and rewards.^5–12^ Though the purpose of such an input region is unclear, theories of the entorhinal cortex have proposed its role in compressing information and encoding it in a format that may be optimal for memory storage.^13,14^ This type of a computation may be a general feature of memory systems.

A challenge to this idea is the fact that many animals – in particular, non-mammals – are adept at hippocampal functions without having an apparent entorhinal cortex. Food-caching birds, for example, rely on the hippocampus to remember locations of many concealed food items in their environment.^15,16^ Recent work has demonstrated mammalian-like firing patterns, including place cells, in the avian hippocampus.^17,18^ Yet birds entirely lack a layered cerebral cortex, which in mammals includes an entorhinal region. Rather than propagating along a cortical sheet, signals in the non-mammalian forebrain are communicated between discrete pallial “nuclei”.^19^ These nuclei are only partially homologous to the mammalian cortex and contain neurons that differ from the cortex in their embryological origins, morphologies, and molecular signatures.^20–23^ It seems unlikely that the avian brain exactly replicates all details of the mammalian microcircuitry relevant for producing characteristic entorhinal activity patterns, like those of grid cells.^24–27^ This raises the possibility than entorhinal-like circuitry and firing patterns are not generally needed for hippocampal function.

Another possibility is that birds have an “interface” region between the hippocampus and the rest of the brain that performs similar functions to the entorhinal cortex. Such a structure may or may not share an evolutionary precursor with the mammalian entorhinal cortex, and it could have substantially different cytoarchitecture and microcircuitry. However, this region would still perform similar computational functions to the mammalian entorhinal cortex. The search for such an entorhinal-like region is hampered by substantial uncertainty and disagreement about the structure of the avian hippocampal formation itself. Using different species, methods, and criteria, studies have proposed more than twenty ways of subdividing the hippocampal formation.^28–33^ The candidate entorhinal cortex in these studies has been placed in at least five different locations, ranging from the midline to the lateral wall of the brain.^28–30,34,35^ Neural recordings in the non-mammalian hippocampal system have so far been relatively scarce, and few have identified activity patterns uniquely characteristic of the entorhinal cortex.^17,36,37^ It is therefore still unknown whether an entorhinal analog exists in the non-mammalian brain.

We set out to address this question using food-caching birds from the *Paridae* family, black-capped chickadees and tufted titmice.^38,39^ Recent work has identified a hippocampal subregion in these species that contains abundant place cells and is likely homologous to the dorsal hippocampus of rodents.^17^ This subregion, as well as the complementary subregion that contains fewer place cells, are well-defined starting points for the anatomical mapping of the circuit. Our goal was to first understand the organization of inputs into the hippocampus. Second, we wanted to understand the physiological similarities, if any, between these inputs and the mammalian entorhinal cortex. Comparing phylogenetically distinct brains in this way could provide powerful insight into the mechanisms of entorhinal function.

## RESULTS

### An input/output region of the avian hippocampus

Entorhinal cortex provides the largest input into the mammalian hippocampus and is a major target of hippocampal output.^3^ It also physically occupies the transitional zone between the hippocampus and the rest of the cortex.^40^ To test whether a similar region exists in birds, we injected retrograde tracers into the hippocampus of a food-caching species, the black-capped chickadee (N=20 injections in 7 birds). Our tracing revealed three inputs into the hippocampus from the avian homologue of the cerebral cortex (the pallium). We first focused on the largest input region, which was immediately lateral to the hippocampus (Figures 1A-B). Hippocampus-projecting cells in this region formed a sharply defined boundary with the hippocampus. These cells were more densely packed than cells in the hippocampus itself, causing the boundary to coincide with a distinct cytoarchitectural transition. We decided to explore this region further because its position adjacent to the hippocampus was similar to that of the mammalian entorhinal cortex.

**Figure 1.**
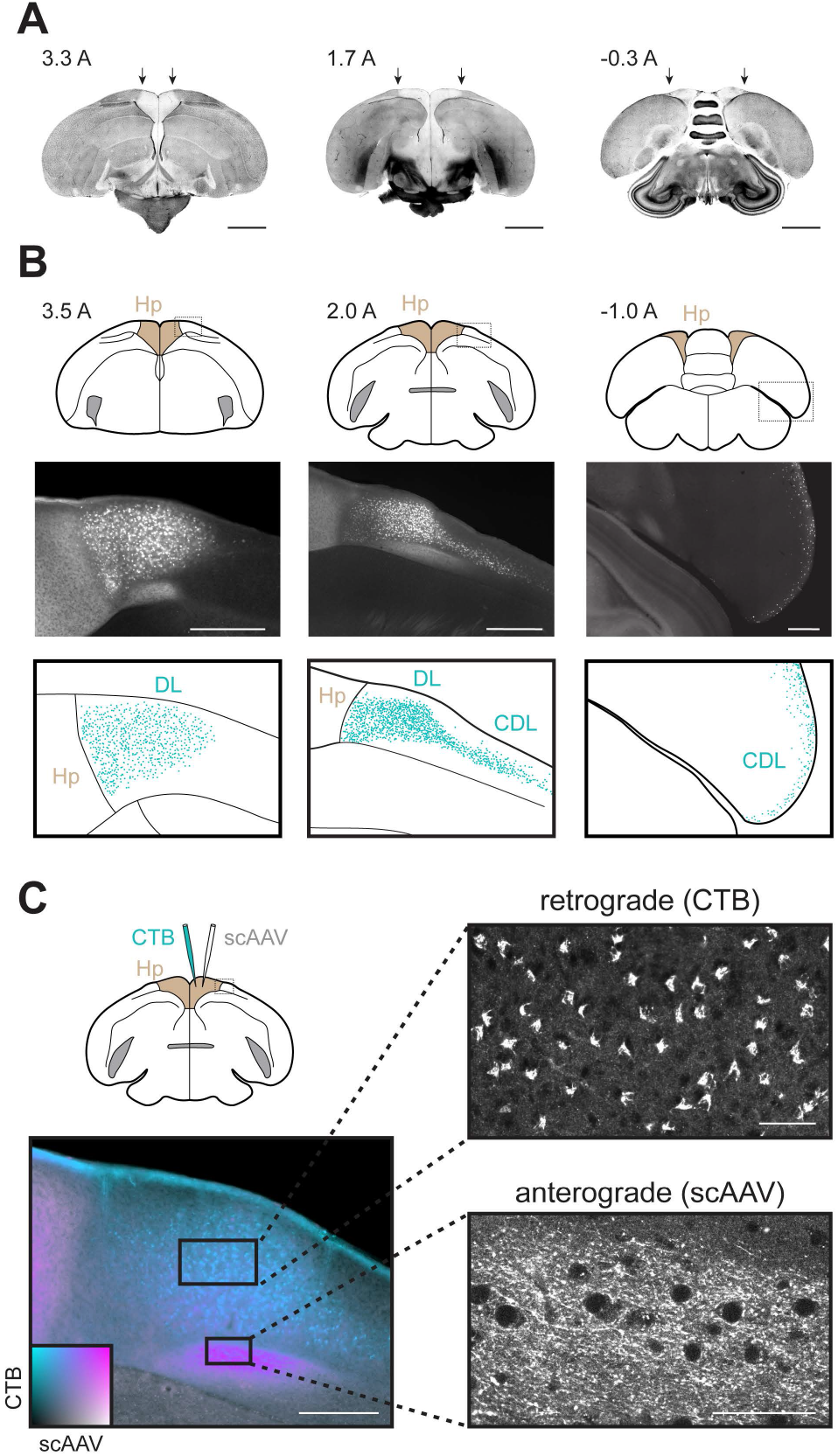
DL/CDL is a major input/output of the chickadee hippocampus. (A) DAPI-labeled coronal slices of the chickadee brain at three locations. Location in mm anterior. Lateral boundaries of the hippocampus are indicated by arrows. The hippocampus is well-delineated cytoarchitecturally and has a lower cell density than the region lateral to it. Scale bars: 2 mm. (B) Retrograde labeling of inputs into the hippocampus. Top: coronal slice outlines at three locations showing the hippocampus (light brown) and the region of interest (dotted rectangle). Middle: fluorescence within the region of interest following CTB injection into the hippocampus. Bottom: labeled cells annotated using a blob detection algorithm. At anterior locations, DL – located adjacent and lateral to the hippocampus – is the most prominent region labeled. In intermediate sections, the transition between DL and CDL is seen. At posterior locations, CDL extends laterally into a thin layer at the surface of the brain. Scale bars: 500 μm. From left to right, injections were at locations 2, 1, and 4 (Table S1). (C) Simultaneous retrograde and anterograde tracing from the hippocampus, using injection of both CTB and a GFP-expressing virus. A two-dimensional color map (inset) was produced by linearly interpolating colors at the four corners, and is used to indicate the amounts of fluorescence produced by the two injections. Axonal fibers are anterogradely labeled by both CTB and GFP and therefore appear in magenta. Cell bodies are retrogradely labeled by CTB only and therefore appear in cyan. The two labels segregate within DL. Scale bar: 250 μm. Higher-magnification confocal images (right) show cell bodies of hippocampus-projecting neurons in dorsal DL and hippocampal axons with axonal terminals in ventral DL. Both injections were at location 1 (Table S1). Scale bars: 50 μm.

Relating this region to known anatomy was nontrivial due to significant discrepancies between published boundaries of the avian hippocampus. Studies relying on cell densities would place our region entirely outside of the hippocampal formation.^41–43^ Other work, using anatomical tracing and forebrain landmarks like the parahippocampal sulcus, would consider it a part of the hippocampal formation.^28,29,44,46^ We decided to use the latter set of criteria, by which the bulk of our region roughly corresponded to the dorsolateral division of the hippocampal formation (DL). The remainder of the region was extended posterior-laterally into a very thin layer, in some places only 1-2 cell bodies thick at the brain surface (Figure 1B). This portion appeared to match the dorsolateral corticoid area (CDL), though it was exceptionally thin in chickadees compared to published species.^33,44^ There was no apparent cytoarchitectural division between DL and CDL; together they formed the largest input into the hippocampus by more than a factor of two, both in volume and in the number of cells.

We asked whether DL/CDL was also a target of hippocampal output. In the mammalian entorhinal cortex, efferent and afferent connections with the hippocampus are segregated between superficial and deep layers, respectively.^4^ We found that a similar pattern of organization was present in the chickadee brain (Figure 1C). Retrogradely labeled somata were positioned dorsally in DL and CDL, while labeled axonal fibers and terminals occupied a deeper part of these structures. We confirmed that these axons belonged to hippocampal neurons by also labeling them with a GFP-expressing virus (scAAV in Figure 1C) injected into the hippocampus (N=4 additional chickadees). Like the entorhinal cortex, DL/CDL is therefore bidirectionally connected with the hippocampus and has a separation of hippocampal inputs and outputs.

### Topographic organization of inputs into the hippocampus

A hallmark of entorhinal anatomy is the topography of its connections with the hippocampus. Different “bands” of the entorhinal cortex project to different sections of the hippocampal long axis, likely contributing to functional differences along this axis.^40,48^ In birds, the apparent equivalent of the long axis is oriented primarily in the anterior-posterior direction.^17,49–51^ To test whether topographic inputs exist in birds, we therefore made injections of retrograde tracers along the anterior-posterior axis of the chickadee hippocampus. We targeted three different sites – either across birds or in the same bird using different fluorophores. We constructed a 3D template model of the chickadee brain and registered all labeled cells to this model.

Retrograde labeling indeed revealed a topographic organization of projections to the hippocampus (Figure 2A). Each hippocampal injection labeled a long, continuous band of cells in DL/CDL. Bands projecting to different sections of the hippocampal long axis were roughly parallel to one another, but curved in a complex way along the surface of the brain, without a consistent orientation relative to stereotaxic axes (Figure 2B). The band of cells projecting to posterior hippocampus included posterior DL and medial CDL. Progressively anterior parts of the hippocampus received inputs from the more anterior parts of DL and more lateral parts of CDL.

**Figure 2.**
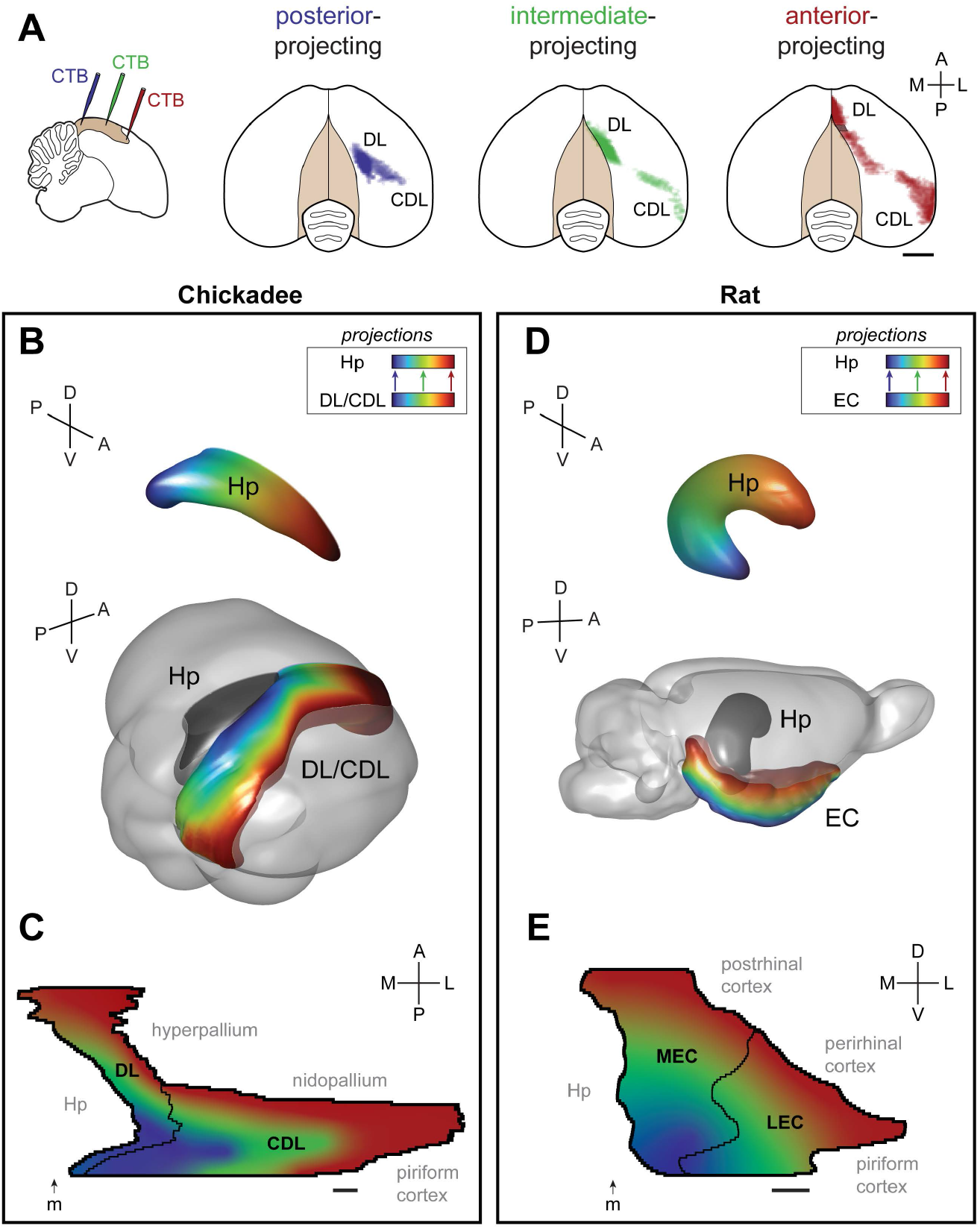
DL/CDL inputs into the hippocampus are topographically organized. (A) Left: outline of a sagittal slice of the chickadee brain, showing locations of three CTB injections into the hippocampus. Right: dorsal views of the brain, showing the density of retrogradely-labeled cells in DL/CDL from each of the injections in one example bird. Each color is applied on a logarithmic scale and individually normalized to its peak fluorescence. Hashed patch shows an area that was too close to an injection site, and where cells were not annotated. Posterior injections label cells in posterior DL and medial CDL. Progressively anterior injections label more anterior DL and more lateral CDL. Scale bar: 2 mm. (B) Bottom: 3D model of a template chickadee brain showing the hippocampus and DL/CDL. Each part of DL/CDL is colored according to the main part of the hippocampus that it projects to, from posterior (blue) to intermediate (green) to anterior (red). Colors are smoothed using a gradient; gradient contours were drawn after registering data like those shown in (A) to the template brain. Top: 3D model of the hippocampus, colored from anterior to posterior using the same color map. (C) Flattened map of DL/CDL formed by unfolding outlines of coronal slices in the chickadee brain. All brain regions bordered by DL/CDL are indicated. Arrow labeled “m” shows the location of the medial ridge, which was used to align outlines of different slices to each other. Map is colored using the same gradient as in (B). Scale bar: 1 mm. (D) Model of the hippocampus and the entorhinal cortex in the rat, constructed using published data (see text). Color gradient is the same as in (B), but indicates location along the dorso-ventral (long) axis of the rat hippocampus. (E) Flattened map of the entorhinal cortex constructed as in (C), but using horizontal slices of the rat brain. Scale bar: 1 mm.

We sought to compare the topographic organization of hippocampal inputs between chickadees and mammals. We noted that all hippocampus-projecting neurons in DL/CDL were near the brain surface (within ∼0.5 mm) and did not have an appreciable topographic organization across depth. This allowed us to treat DL/CDL as a two-dimensional structure and construct its flattened map using classic cortical “unfolding” techniques that are usually not possible in the nucleated avian brain.^40,52^ Figure 2C shows a map constructed by unfolding outlines of annotated coronal slices of the chickadee brain.

Compared to its avian homolog, the rodent hippocampus appears to be rotated in stereotaxic coordinates by approximately a 90-degree “backward pitch” (Figure 2D). Because of this rotation, the anterior-posterior axis of the bird hippocampus is likely equivalent to the dorso-ventral axis in rodents.^17,49–51^ It occurred to us that the same transformation may extend to neighboring entorhinal-like regions. In rodents, the entorhinal cortex borders the hippocampus medially on the posterior surface of the brain, then extends to the lateral and ventral surfaces. In birds, DL/CDL also borders the hippocampus medially, but on the dorsal brain surface (Figure 2B). It then extends onto the lateral and posterior surfaces. To compare topographic maps constructed from avian coronal slices, we therefore used horizontal slices of the rat brain, which are related to coronal slices by a 90-degree pitch rotation. Figure 2E shows such a map constructed using published data.^40,53–55^

Flattened maps of the chickadee DL/CDL and the rat entorhinal cortex were remarkably similar. Both regions bordered the hippocampus medially and cortical structures laterally. In rats, cortical structures included the piriform cortex and neocortical areas (postrhinal and perirhinal cortices). In chickadees, structures in the same relative positions also included the piriform cortex, as well as hyperpallium and nidopallium. In both species, topographically organized bands of hippocampal projections were oriented orthogonally to the border with the hippocampus. Therefore DL/CDL and the entorhinal cortex were oriented similarly relative to other brain areas and had topologically equivalent organizations of hippocampal connections. Interestingly, the relative positions of medial and lateral entorhinal cortices (MEC and LEC) were similar to that of DL and CDL, respectively, suggesting a possible analogy of these regions.

### Pathways for cortical inputs into the hippocampus

We next asked what role DL/CDL played in relaying cortical inputs into the hippocampus. In mammals, many cortical regions project to the entorhinal cortex, but only a small subset of these project directly to the hippocampus.^3^ We sought to perform the same comparison in birds. We focused on DL, which is by far the larger part of the input structure in chickadees and could be targeted using conventional retrograde tracing techniques.

Direct inputs into the hippocampus, besides DL/CDL, included two cortical structures, consistent with those described in other avian species ^28,35,56^. One of these was hyperpallium densocellulare (HD; Figure 3A). This region is primarily the target of the thalamofugal visual pathway, though it might also integrate other sensory modalities.^57,58^ The other input originated in the lateral part of the nidopallium (NL), including its frontal (NFL) and intermediate (NIL) portions (Figure 3B). This region is likely the target of the tectofugal visual pathway.^59^

**Figure 3.**
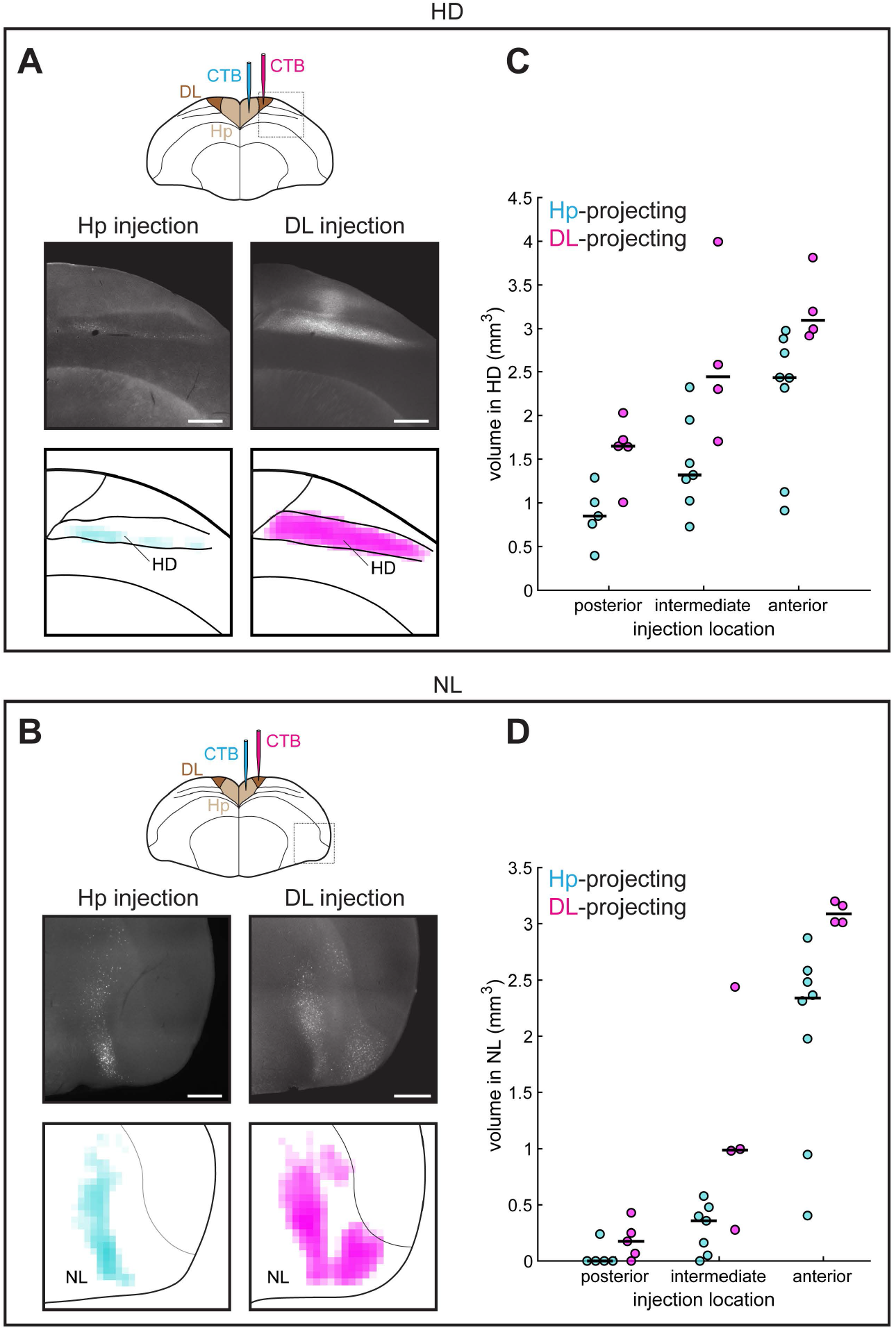
DL/CDL is the main target of cortical pathways into the hippocampus. (A) Top: outline of a coronal slice of the chickadee brain, showing injections of CTB into the hippocampus and into DL. Middle: examples of retrograde tracing in HD. Bottom: densities of retrogradely-labeled cells in HD in the same example images. Each color is applied on a logarithmic scale and normalized to the maximum across both plots. Whereas hippocampal injections label only the ventral part of HD, DL injections label the entire dorso-ventral extent of HD. Scale bar: 1 mm. Injections were at locations 1 and 5 (Table S1). (B) Retrograde labeling in NL, shown as in (A). Injections were at locations 4 and 7 (Table S1). (C) Volume of tissue in HD retrogradely labeled by injections into the hippocampus and into DL. Each symbol is an individual injection. Horizontal lines indicate medians. At each location, more volume in HD was labeled by DL injections than by hippocampal injections. (D) Quantification of retrograde labeling in NL, shown as in (B).

We next injected retrograde tracers into DL (N=13 injections in 9 birds). We found that cortical inputs into DL originated from the same two structures, HD and NL (Figures 3A-B). However, in both cases, these inputs included additional portions of the structure that did not project directly to the hippocampus. In HD, hippocampus-projecting neurons were limited to the ventral portion of the region, whereas DL-projecting neurons occupied the entire dorso-ventral extent (Figure 3A). In NL, the DL-projecting cells occupied an additional lateral portion that only sparsely projected to the hippocampus (Figure 3B).

To quantify these observations, we compared the volume of tissue retrogradely labeled by injections into the hippocampus and into topographically matched locations in DL. We considered each of the three anterior-posterior locations separately because in both HD and NL there were vastly more neurons projecting to anterior locations than to posterior locations (p<0.005 regression of volume by anterior position). In HD, the volume of tissue labeled by DL injections compared to hippocampal injections was larger in all three cases (p<0.05 Wilcoxon rank-sum test, Figure 3C). In NL, this difference was significant (p<0.05) for all injections except the most posterior one, which barely labeled any cells (Figure 3D). Our results show that DL/CDL, like the entorhinal cortex, is the primary target of the cortical pathway into the hippocampal formation.

### Spatial activity in DL

Our anatomical analysis led to clear predictions about neural activity in DL/CDL. In particular, the anterior portion of DL in our mapping appeared to be equivalent to the dorsal MEC of rodents^40^ – the subregion renowned for its spatially selective firing.^10,60^ We asked whether activity patterns in these regions were indeed similar. To record DL neurons, we adapted one-photon calcium imaging^61^ for freely-behaving tufted titmice – a species from the same family as chickadees that is larger and therefore more amenable to head-mounted microscopes (Figures 4A-B). We trained titmice to forage for scattered seeds in an environment comparably sized to those in which periodic firing patterns were observed in rodents.^10^ Our approach was to first analyze DL activity for specific types of patterns that have been described in the dorsal MEC. Afterwards, we fit activity with a less biased statistical model that allowed for complex types of selectivity.

**Figure 4.**
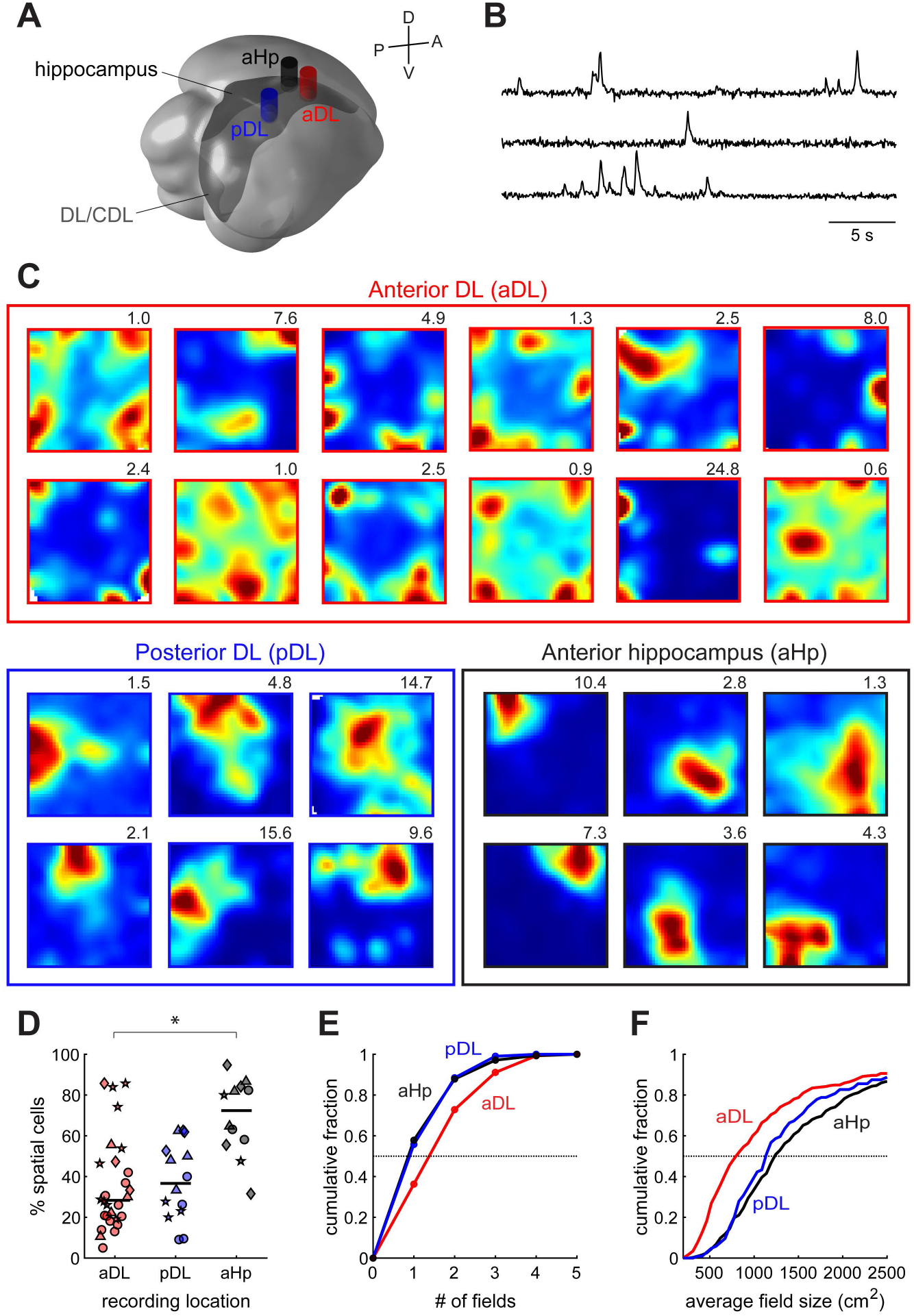
Cells in anterior DL exhibit multi-field patterns of spatial activity. (A) Locations of three imaging coordinates in the titmouse brain. Each cylinder indicates the position and extent of the GRIN lens on the brain surface. (B) Example calcium traces simultaneously recorded in anterior DL using a head-mounted one-photon microscope. Fluorescence is in arbitrary units extracted by the CNMF_E algorithm.^94^ (C) Examples of spatial maps from cells recorded in each of the three regions. For each cell, color map ranges from 0 (blue) to 99th percentile of the pixels (red). Maximum firing rate in calcium events per second is shown above each map. Most spatial cells in anterior DL exhibit multiple firing fields, whereas typical cells in posterior DL and the hippocampus exhibit one larger firing field. Examples are intentionally chosen to show multi-field cells we discovered in anterior DL and illustrate the main difference between the regions; an unbiased selection of firing patterns is shown in Figure S1. (D) Fraction of cells identified that were significantly spatial in each of the recording sessions. Different symbols are used for different birds. Horizontal lines indicate medians. Spatial cells were more prevalent in the anterior hippocampus (p<0.001 Wilcoxon rank-sum test), but were abundant in all three recorded regions. (E) Cumulative histograms of firing field size for spatial cells in each of the recorded regions. (F) Cumulative histograms of the number of firing fields per spatial cell in each of the recorded regions.

A total of 1270 cells were recorded in the anterior DL of four birds. Because this number is likely to double-count some of the same cells recorded across days, we verified all of our results with analyses that first pooled data for each bird. A large fraction of the cells in anterior DL were spatial (39±8% across birds, mean±SEM here and elsewhere, Figures 4C-D, S1). Most spatial cells displayed multi-field firing maps that were unlike those previously described in birds.^17^ Of the 405 spatial cells, 258 (64%) had more than one firing field, with many cells having three or more fields (Figure 4E). These cells superficially resembled grid cells, but their fields were not arranged in a periodic lattice (none of the cells passed the standard test for “gridness”.)^62^ This irregular distribution of firing fields resembled MEC activity in three-dimensional tasks,^47,63^ even though our birds moved primarily in two dimensions during the experiment. Like in flying bats,^47^ firing fields of some cells were spaced more regularly than expected by chance – suggesting that some local organization was present in DL activity (Figure S2). By analogy with this study, we refer to these cells as “grid-like” even though their firing fields do not conform to a grid.

Are multi-field firing patterns unique to DL, or do they result from some of our experimental parameters, like the use of a large environment?^64–66^ To test this, we recorded 463 cells in the anterior hippocampus in four additional titmice. As in prior work on titmice,^17^ the hippocampus in our recordings exhibited abundant spatial activity (69±3% of cells across birds, Figures 4C-D, S1). Although some of these cells had multiple fields, they on average had fewer fields than cells in DL (p<0.001 Chi-squared test, Figure 4E). In contrast to DL neurons, most hippocampal spatial cells had only one firing field (58%). Fields in the hippocampus were also larger than the fields in DL (p<0.001 Wilcoxon rank-sum test, Figure 4F). Therefore, multi-field firing patterns are not ubiquitous in the hippocampal circuit, but may be more typical of DL.

Next, we asked whether spatial representations differed across the anterior-posterior axis of DL. MEC in rodents has a dorso-ventral organization that parallels its topographic connections with the hippocampus.^10,40,67,68^ Whereas dorsal MEC has finer spatial coding, with neurons exhibiting larger numbers of smaller firing fields, spatial coding in ventral MEC is coarser. We recorded 274 neurons in the posterior part of DL in four additional titmice. Many of the neurons in this region were spatial (38±9% across birds, Figures 4C-D, S1), but exhibited larger firing fields than neurons in anterior DL (Figure 4F, p<0.001 Wilcoxon rank-sum test). Compared to anterior DL, these cells had fewer fields, with most cells having only one (p<0.001 Chi-squared test, Figure 4E). The anterior-posterior axis of DL therefore seems to be organized functionally, much like its mammalian counterpart in MEC.

Another canonical feature of spatial activity the mammalian MEC are border cells – neurons that fire near one or multiple boundaries of the environment.^60^ In anterior DL we also observed abundant cells with firing near the walls of the arena (Figure 5A). Of the 405 spatial cells, 83 (21%) were considered border cells by the standard analyses used in mammals. Unlike in anterior DL, border cells were nearly absent in the anterior hippocampus (4% of the 313 spatial cells) and posterior DL (1% of the 104 spatial cells, Figure 5B). Border cells, like multi-field activity, may therefore also be a specialization of anterior DL.

**Figure 5.**
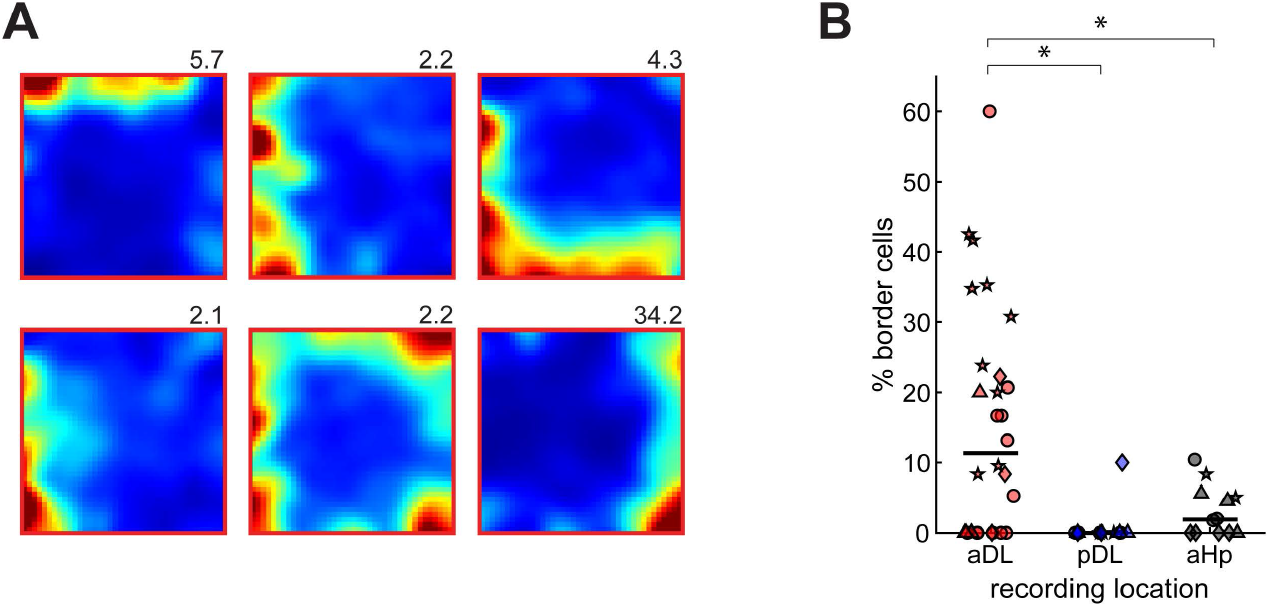
Anterior DL has abundant border cells. (A) Spatial maps of example border cells in anterior DL, plotted as in Fig. 4. (B) Fraction of spatial cells identified as border cells in each of the recording sessions. Different symbols are used for different birds. Border cells were more abundant in anterior DL than in posterior DL (p<0.001, Wilcoxon rank-sum test) and anterior hippocampus (p<0.05).

Finally, we examined the temporal characteristics of these spatial neurons. In all species tested, including birds, hippocampal spatial activity is “prospective”: it is most strongly correlated to the animal’s position in the future.^17,69–71^ In MEC, however, activity is less prospective, with latencies closer to zero.^72,73^ We asked whether a similar pattern existed in birds. We measured spatial information encoded by each cell’s activity at various lags relative to the behavioral trajectory and determined the lag at which spatial information was maximized (Figure 6A). As in previous studies, most spatial cells in the hippocampus had a positive (prospective) lag with an average of 350±40 ms (p<0.001 t-test, N=313 cells, Figure 6B). In contrast, the lag was close to zero in DL, and even significantly negative (retrospective) in posterior DL (−40±40 ms and -400±100 ms in anterior and posterior DL, respectively, N=405 and 104 cells, p<0.001 comparison to the hippocampus in both cases). These analyses show that hippocampal inputs from MEC in mammals and from DL in birds are not only spatially similar, but have analogous temporal characteristics.

**Figure 6.**
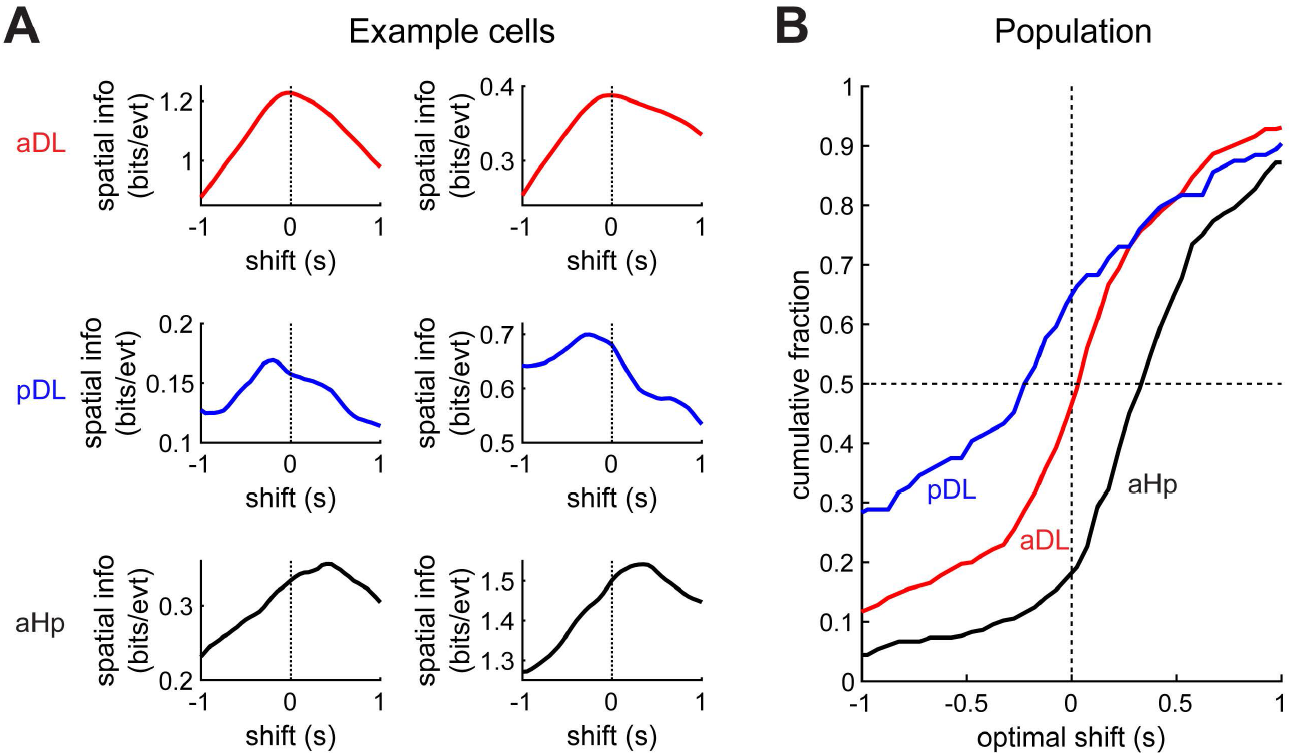
DL, like the entorhinal cortex, is less prospective than the hippocampus. (A) Examples of six cells in the three recorded titmouse brain regions. For each cell, spatial information is quantified at different temporal shifts of spikes relative to the behavioral trajectory. The optimal shift tends to the positive (prospective) for the hippocampus, close to zero for anterior DL, and retrospective (negative) for posterior DL. (B) Cumulative histograms of the optimal shifts for cells in each of the recorded regions.

### Representation of other navigational variables in DL

Mammalian MEC encodes an array of navigational variables other than position – notably the animal’s head direction and speed.^11,12^ We found both of these variables represented in the activity of titmouse DL neurons. In anterior DL, 34±5% of cells across four birds had significant head direction tuning (Figures 7A-B), and 40±9% of cells had significant speed tuning (Figures 7C-D). Speed cells included those with positive and those with negative correlations between speed and firing rate, though the former were more common (77%, p<0.001 Chi-squared test). Consistent with recordings along the dorso-ventral axis of MEC,^74^ we found both head direction and speed cells in both anterior and posterior DL (Figures 7B,D). As in previous studies, these types of cells were also found in the hippocampus (Figures 7B,D).

**Figure 7.**
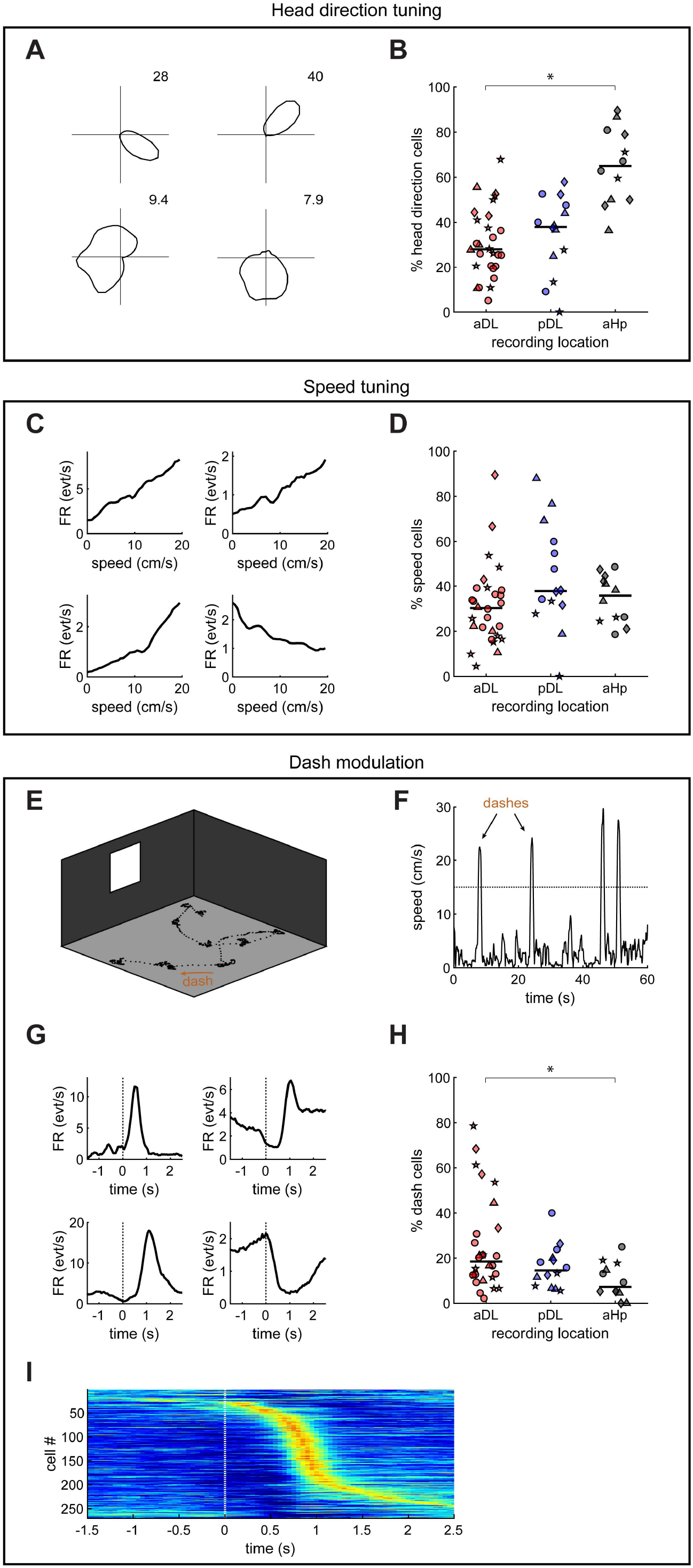
DL represents a variety of navigational variables other than location. (A) Examples of head direction cells in anterior DL, showing two cells with narrow tuning (top) and two cells with broader tuning (bottom). Each plot shows firing rate as a function of head direction in polar coordinates. Peak firing rate in calcium events per second is indicated for each cell. (B) Fraction of cells identified as head direction cells in each of the recording sessions. Different symbols are used for different birds. Horizontal lines indicate medians. Head direction cells are found in all three regions, with higher abundance in the hippocampus (p<0.001 Wilcoxon rank-sum test). (C) Examples of speed cells in anterior DL, showing three cells with positive speed tuning and one with negative speed tuning. (D) Fraction of cells identified as speed cells, plotted as in (B). Speed cells were found in all three regions. (E) Example 3 min trajectory of the titmouse overlaid on a 3D schematic of the recorded arena. Behavioral data points are recorded every 50 ms. Trajectory indicates periods of relative immobility separated by fast, ballistic “dashes”. (F) Example recording of the bird’s speed, showing four dashes. (G) Examples of cells in anterior DL with modulation by time relative to the dash. Zero indicates dash onset. Each trace is an average across all dashes recorded in the session. (H) Fraction of cells identified as dash-modulated cells, plotted as in (B). Dash-modulated cells were found in all three regions, but were more abundant in anterior DL compared to the anterior hippocampus (p<0.05 Wilcoxon rank-sum test). (I) Activity of all dash-modulated cells in anterior DL, sorted according to the time of the peak firing rate. Each cell’s activity is normalized from 2.5^th^ (blue) to 97.5^th^ (red) percentile of its firing rates in the plotted window. Firing of all cells is concentrated around 0.5-1 s relative to the dash, but forms a sequence spanning the entire event.

Avian behavior offers an additional opportunity to study how entorhinal-like activity represents the animal’s movement. Titmice, like most small passerine birds, move on the ground by hopping rather than walking. In a foraging task, they approach a food target via a ballistic sequence of hops (“a dash”) lasting about 1 s (Figures 7E-F). We found that activity of many cells in anterior DL was modulated by these dashes (30±8% across birds, Figures 7G-H). Much of the activity was concentrated between 0.5 and 1.5 s after the onset of the dash, roughly corresponding to the time birds were approaching the target. However, different cells had different latencies, forming a sequence of activity that spanned the entire dash (Figure 7I). These differences in latency were consistent across trials within a session were therefore not due to noise (p<0.001, shuffle test). This activity also could not be explained entirely by speed tuning: for example, cells that fired at the end of the dash did not generally fire during similar speeds at the beginning of the dash. We found some dash tuning in the posterior DL and the anterior hippocampus as well, but it was weaker and involved fewer cells than in the anterior DL (Figure 7H). Altogether, our results so far show that DL represents a remarkable diversity of behaviorally-relevant variables in a spatial task.

### Mixed selectivity of DL neurons

How are these different encoded variables organized across the DL population? For each pair of variables we considered (location, head direction, speed, and time relative to a dash), we found DL cells representing both variables, only one variable, or neither (Figure 8A). Across the population, there was no evidence of distinct classes of cells encoding individual variables. These results suggest that DL might recapitulate an important property of the entorhinal cortex and mammalian cortex in general – the mixed selectivity of cells to task-related parameters.^11,45,75,76^

**Figure 8.**
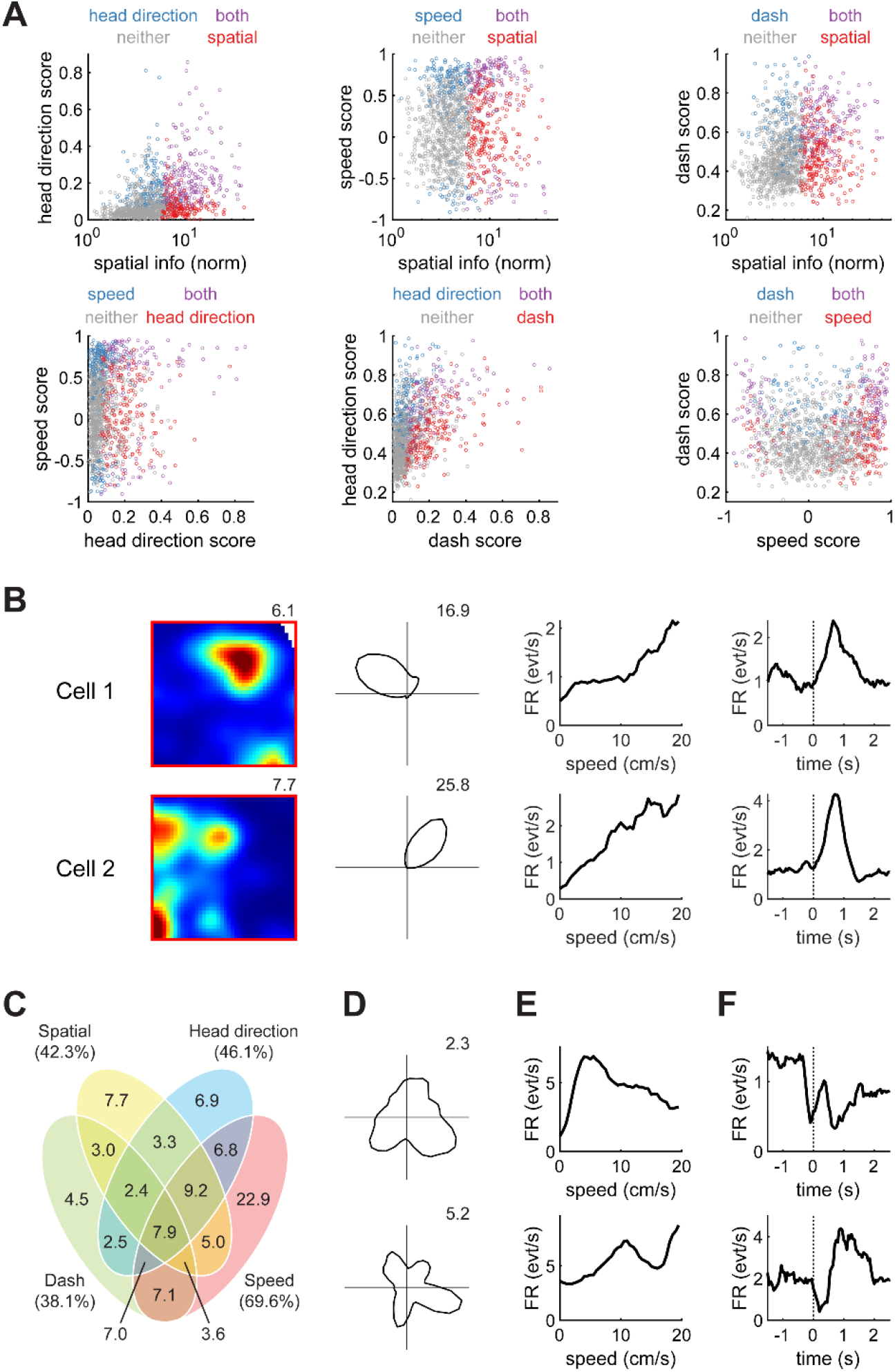
DL exhibits mixed representation of navigational variables. (A) Representation of pairs of variables by anterior DL neurons. For each pair of variables (location, head direction, speed, and time relative to dash), each cell’s selectivity to those variables is shown. For each pair, there are cells representing neither variable, representing only one variable, or representing both. (B) Examples of two cells identified by the linear-nonlinear-Poisson model as being modulated by all four analyzed variables. Conventional tuning curves are plotted, as in Figures 4 and 7. (C) Venn diagram of anterior DL cells, classified according to what subset of the four variables they were modulated by. The diagram classifies 1002 cells that were considered by the model to be modulated by at least one variable; an additional 268 cells were not modulated by any variable. (D) Examples of head direction tuning curves showing multiple peaks. These cells were not detected as head direction cells by standard analysis, but were identified as head direction-modulated by the model. (E) Examples of non-motonotic speed tuning identified by the model. (F) Examples of complex modulation by time relative to dashes identified by the model.

So far, we have focused on searching for specific, hand-picked types of neurons inspired by the mammalian literature. We next sought to use a less biased approach to characterize DL activity. First of all, we wanted to account for complex tuning curves – e.g., tuning of a cell to multiple head directions, non-monotonic speed tuning, etc.

Second of all, we wanted to account for possible conjunctive tuning of individual cells to multiple variables, not only pairs of variables. Finally, we wanted to account for arbitrary correlations between these variables. For example, due to biases in an animal’s trajectory, a place-selective cell might appear to be tuned to head direction, or a speed-tuned cell might appear to be modulated by dashes. We implemented a linear-nonlinear-Poisson model based on one developed for rodent MEC.^45^ This model fits each cell’s activity as a linear combination of responses to multiple behavioral variables. The tuning curve for each variable is allowed to be an arbitrarily complex shape. Using cross-validation to prevent overfitting, the model selects, for each cell, the subset of variables that improve the model fit and therefore contribute to that cell’s firing.

The model detected 79% of the 1270 cells in anterior DL as selective to at least one of the behavioral variables we considered. The remaining minority of cells were best fit by a constant firing rate fit to the entire behavioral session (but, of course, could be selective for some variables we did not measure). Many cells were best fit by a combination of two, three, or four variables, with every possible subset of these variables present in the recorded population (Figure 8B-C). In some cases tuning curves exhibited complex shapes that were not accounted for by the statistical measures we previously considered: for example, some cells had non-monotonic tuning for speed, while others had multi-peaked modulation by head direction and dashes (Figures 8D-F). Similar types of complex tuning and mixed selectivity were also found in the posterior DL and the hippocampus (Figure S3). Overall, our results demonstrate highly complex, mixed representations of navigational variables in the hippocampal system of food-caching birds. Much like in the mammalian MEC, these multiplexed representations are already present in DL – the major input into the hippocampus.

## DISCUSSION

We have identified a brain region (DL/CDL) in food-caching birds that has many anatomical and physiological similarities to the mammalian entorhinal cortex. Anatomically, this region is located at the interface between the hippocampus and the rest of the cortex, and appears to be the main route of information in and out of the hippocampus. Like the entorhinal cortex, it is topographically connected with the long axis of the hippocampus and parallels the hippocampal organization of activity along this axis. Physiologically, DL contains most cell types identified in rodents during foraging tasks – including multi-field cells, border cells, head direction cells, speed cells, and neurons with complex conjunctive representations. These similarities are present in spite of birds being highly divergent from mammals in brain development and architecture, most notably lacking a layered cortex. An avian structure with so many characteristic features of the entorhinal cortex was therefore unexpected, and emphasizes the fundamental role these features likely play in hippocampal computation.

In addition to analogous spatial representations in the mammalian MEC and avian DL, we found an interesting similarity in the timing of these signals. Whereas the hippocampi of both mammals and birds are predictive of the animal’s future location, their inputs from both MEC and DL are much less predictive.^17,69–73^ Converting incoming signals into a prediction of the future may therefore be a function of the hippocampus itself, and this function might be conserved across vertebrates. This computational role of the hippocampus is consistent with theoretical work on predictive representations.^13^

A notable difference between DL and the mammalian entorhinal cortex is the lack of cells with regularly arranged grid firing fields. The firing we observed in DL instead resembles irregular patterns reported in MEC during 3D navigation.^47,63^ One of these studies has proposed that switching between regular and irregular arrangements of fields can be achieved by changing a single “temperature” parameter in a spatial system.^47^ Perhaps avian DL is in a parameter regime that produces irregular arrangements of fields even during 2D navigation. One way this could occur is via differences in the underlying microcircuit. For example, grid patterns have been proposed to depend on the precisely tuned connections between excitatory and inhibitory neurons,^24–27^ which are likely divergent across phylogenetically distant species.^20–22^ Alternatively, some models have suggested that field arrangement can be influenced by the statistics of movement in the environment,^13^ which is also different between hopping birds and walking mammals. Our results add to the growing body of evidence that, even though multi-field firing patterns are fundamental to the entorhinal cortex, their regularity is not a requirement for the function of this region.

Several previous studies have presented evidence linking DL to the entorhinal cortex. For instance, anatomical tracing and electrical stimulation experiments in pigeons have suggested a general pattern of information flow in the lateral-to-medial direction within the hippocampal formation.^35,56,77^ Some authors have proposed that the lateral parts of the circuit are therefore analogous to the entorhinal cortex.^28,46,56^ However, a sharp delineation of input/output subregions in the hippocampal formation has not been found in other species so far, and might be unique to chickadees. The entire hippocampal circuit is larger in chickadees and other food-caching birds, so it is possible that a specialization of distinct subregions has accompanied this evolutionary enlargement. Such specialization might be a common phenomenon across other enlarged neural systems, such as the visual system in primates and the song system in songbirds.^78,79^

Unlike DL, CDL – the lateral part of the region we identified – has not been generally proposed as an entorhinal analog in previous studies. According to our anatomical analysis, CDL seems to occupy the same position as the rodent LEC (Figure 2). In rodents, LEC borders the piriform cortex and is the primary source of olfactory inputs into the hippocampus,^80,81^ though it also processes other sensory modalities.^82,83^ Notably, CDL in birds is also adjacent to the piriform cortex and in other species receives inputs from the olfactory system.^44,84^ This role might explain its exceedingly small size in chickadees, which barely use smell and have one of the smallest olfactory bulbs among vertebrates.^85^ It remains to be seen whether CDL has functional similarities to LEC. Neural recordings of this region might be more practical in other avian species.

What accounts for the presence of an entorhinal-like region in birds? There are at least three explanations, which are not mutually exclusive. One possibility is that DL/CDL and the entorhinal cortex are derived from the same evolutionary precursor and have conserved some of the same ancestral functions. Our topological analysis places DL/CDL in the same location as the entorhinal cortex relative to other brain regions – at the lateral terminus of the hippocampal transverse axis. This axis shares multiple gene expression signatures between mammals and non-mammals,^21,86^ suggesting that DL/CDL and the entorhinal cortex might also be transcriptionally similar. It is conceivable that even partially conserved cells types and connectivity are sufficient to account for the entorhinal-like activity patterns we observed. Of course, there are also likely to be significant transcriptional differences between DL/CDL and the entorhinal cortex. DL/CDL might also combine the genetic features of multiple adjacent mammalian regions – much like a region recently described in amphibians that combines entorhinal and subicular cell types.^22^ Detailed transcriptomics data in birds will hopefully shed light on these possibilities.

A second possibility is that entorhinal-like activity in birds is the result of convergent evolution. A large body of theoretical work has identified the benefits of entorhinal-like activity patterns in a spatial memory system. For example, multi-field representations with different field spacings across cells are a particularly efficient way to encode spatial location by neural activity.^14,66,87^ Activity of head direction cells and border cells is thought to be indispensable for the spatial computations in the circuit.^11,60,88^ Work across brain regions has also shown how complex forms of mixed selectivity, like the ones observed in DL/CDL and the entorhinal cortex, can increase the computational capacity and flexibility of a neural network.^11,45,75,89^ It is possible that evolutionary pressure has forced even unrelated neural circuits to produce similar entorhinal-like patterns of activity. This pressure might be particularly intense in food-caching birds, which use their hippocampal system for storing and recalling exceptionally large numbers of spatial memories.^15,38,90^

The third possibility is that some entorhinal-like patterns of activity arise naturally within the hippocampal circuit. For instance, recent work has shown that grid cells can appear in neural networks that are simply trained to perform certain spatial navigation tasks.^91,92^ This process might not require highly specific circuitry, but could occur in an arbitrary recurrent network exposed to appropriate behavioral demands. Other work has shown that grid-like firing can result from dimensionality reduction operations on the activity of place cells.^13^ These types of operations could be implemented naturally by a network in which a smaller number of neurons receive feedback from a larger population of hippocampal neurons^93^ – a general architecture that appears to characterize both the avian DL/CDL and the mammalian entorhinal cortex.

Studies in mammals have focused on the roles of specific cell types in generating entorhinal activity. They have also begun to characterize in detail the microcircuit organization, connectivity, and plasticity rules within this brain area. Although some of these details may be similar between birds and mammals, many are likely to be quite different. The discovery of an entorhinal-like region in birds allows for a potentially powerful comparative approach to study each of these features. It may indicate which features of entorhinal anatomy and physiology are fundamental to entorhinal function, and which ones are specializations unique to particular species of animals.

## ACKNOWLEDGEMENTS

We thank Stephanie Hale, Selmaan Chettih, Felix Moll, Michael Long, and Luke Hammond for technical assistance; the Black Rock Forest Consortium, Timothy Green, and Jennifer Scribner, for field site help; Selmaan Chettih, Isabel Low, and Hannah Payne for comments on the manuscript. Imaging was performed with support from the Zuckerman Institute’s Cellular Imaging platform. This work was supported by the Beckman Foundation Young Investigator Award, the New York Stem Cell Foundation — Robertson Neuroscience Investigator Award, NIH Director’s New Innovator Award (DP2-AG071918), NIH training grant (T32 EY013933, to MCA), NSF Graduate Research Fellowship Program (to MCA), Simons Society of Fellows (to ELM).

## AUTHOR CONTRIBUTIONS

M.C.A and D.A conceived of the presented idea. M.C.A, K.S.G, and E.L.M performed the experiments with supervision from D.A. E.L.M wrote code needed for calcium imaging analyses. M.C.A and D.A analyzed the data and wrote the manuscript.

## METHODS

### Animal subjects

All animal procedures were approved by the Columbia University Institutional Animal Care and Use Committee and carried out in accordance with the US National Institutes of Health guidelines. The subjects were 17 black-capped chickadees (*Poecile atricapillus*) and 12 tufted titmice (*Baeolophus bicolor*) collected from multiple sites in New York State using Federal and State scientific collection licenses. Subjects were at least 4 months old at the time of the experiment, but age was not determined more precisely.

Neither chickadees nor titmice are visibly sexually dimorphic. All surgical and behavioral procedures were performed blindly to sex, and sex was determined only post-mortem after the experiments. Of the chickadees, 13 (4 female, 9 male) were used for retrograde tracing, 4 (all male) were used for anterograde tracing, and one (male) was used to create a 3D rendering of the brain. All titmice (7 female, 5 male) were used for awake behaving calcium imaging experiments.

Prior to experiments, all birds were housed in an aviary in groups of 1-3, on a ‘winter’ light cycle (9 h:15 h light:dark). Birds were given an ad-libitum supply of a small-bird diet (either Small Bird Maintenance Diet, 0001452, Mazuri or Organic High Potency Fine Bird foods, HPF, Harrison’s Bird Foods). Upon transfer to the lab for experiments, birds had primary flight feathers trimmed and were individually housed.

### Surgery

For all surgical procedures, birds were anesthetized with 1-2% isoflurane in oxygen. Feathers were removed from the surgical site, and the skin was treated with Betadine. Birds were then placed into a stereotaxic apparatus for the duration of the surgery. The head was rotated to an angle appropriate for each experiment (see below), measured as the angle between the groove in the bone at the base of the upper beak mandible and the horizontal table surface. Throughout the surgery, saline was injected subcutaneously to prevent dehydration (∼0.4 mL every 30-60 min). After the surgery, an analgesic (buprenorphine, 0.05 mg/kg) was injected intraperitoneally.

#### 3D brain reconstruction

To facilitate surgical planning and visualize the 3D arrangement of brain structures, we constructed a template chickadee brain. The template included boundaries of the hippocampus, DL/CDL, and the brain surface. In one bird, stiff wire (Malin Co., 0.006” music wire) was inserted horizontally from the lateral wall of the brain toward the midline to create two visible tracts. The bird was immediately perfused (see below), and following tissue fixation, the brain was sliced sagittally. Slices were annotated (Adobe Illustrator) and aligned using custom MATLAB code, with the two lesions used for alignment.

#### Anatomical tracing experiments

For anatomical tracing, all injections were made in chickadees at a 75° head angle. Injection coordinates for different brain regions are given in Table S1. For retrograde tracing, we used cholera toxin subunit B (CTB in 1X PBS, 0.4% weight/volume) fluorescently conjugated to Alexa Fluor in one of three different wavelengths (488 nm, 555 nm, and 647 nm; C34775, C34776, C34778, respectively, Invitrogen). We made injections in 1-3 tracing locations per animal, using different wavelengths and sometimes different hemispheres for different locations in individual birds. For the anterograde tracing experiment, scAAV9-CBh-GFP virus (UNC viral core) was injected in the posterior hippocampus.

**Table S1:** Stereotaxic coordinates for the center of injection locations in anatomical tracing and calcium imaging experiments. *All coordinates are in mm. AP and ML coordinates and relative to lambda. DV coordinates are relative to the brain surface. A 75*° *head angle was used for chickadees and a 65*° *head angle for titmice. The location listed as* “*Figure 1B, left*” *was used only for illustration purposes in that figure, but not any of the analyses*.

| Location index | Species | Region | Sub-region | AP coord. | ML coord. | DV coord. |
| --- | --- | --- | --- | --- | --- | --- |
| 1 | Chickadee | Hippocampus | Posterior | 1.0 | 0.4 | 0.4 |
| 2 |  |  | Figure 1B, left | 2.0 | 0.4 | 0.4 |
| 3 |  |  | Intermediate | 3.0 | 0.4 | 0.4 |
| 4 |  |  | Anterior | 5.0 | 0.4 | 1.0 |
| 5 |  | DL | Posterior | 2.0 | 2.3 | 0.4 |
| 6 |  |  | Intermediate | 3.5 | 1.7 | 0.4 |
| 7 |  |  | Anterior | 5.5 | 0.7 | 0.3 |
| 8 | Titmouse | Hippocampus | Anterior | 3.5 | 0.7 | 0.4 |
| 9 |  | DL | Posterior | 2.0 | 2.6 | 0.4 |
| 10 |  |  | Anterior | 4.5 | 1.4 | 0.4 |

Injections were made using Nanoject II (Drummond Scientific) and a pulled glass pipette. Immediately after the pipette was lowered into the brain, the brain surface around the pipette was sealed with a silicone adhesive (Kwikcast, World Precision Instruments). This prevented the surface from drying out and injected liquid from leaking out of the brain. For retrograde tracing, 2 injections of 50 nL of CTB were made over the course of 5 min at each location. For anterograde tracing, one 385 nL injection of the virus was made over the course of 7 min. In both cases, the pipette was left in place for an additional 5 min after the injections. For all anatomy experiments, the animal was perfused one week following surgery.

#### Surgeries for calcium imaging

For calcium imaging, all injections were made in titmice at a 65° head angle. At the beginning of the surgery, dexamethasone (2 mg/kg) was injected intraperitoneally to prevent inflammation around the implant. All recordings were conducted in the right hemisphere. To anchor the implant, 6 partial craniotomies were made through the top layer of the skull, surrounding the recording location. Insect pins were inserted to connect pairs of these craniotomies to each other. These insect pins and the craniotomies were then covered with light-cured dental cement (D69-0047, Pearson Dental), fully surrounding the recording location. A full craniotomy and durotomy of ∼1.4 mm diameter were then made over the recording location.

For all experiments, viral injections were used to induce GCaMP6f expression (AAV9-CAG-GCaMP6f-WPRE-SV40, 100836-AAV9, Addgene). For the hippocampal recordings, an injection of 880 nL of the virus was made at over the course of 11 min, and then the pipette was left for an additional 25 min. For the DL recordings, a grid of 12 injections spaced 0.2 mm apart was made into DL coordinates. Each DL injection contained 58 nL administered over 3.5 min, and the pipette was left in the brain for an additional 3 min. Clear silicone elastomer (Kwiksil, World Precision Instruments) was used to seal the brain surface during all virus injections.

Following the injections, for the hippocampal recordings, a 1 mm diameter GRIN lens (1050-004595, Inscopix) was lowered 0.3 mm into the brain, and the surrounding exposed brain surface was covered with the clear elastomer. The lens was then secured to the outer anchor points using light-cured dental cement.

For the DL recordings, a thin layer of the clear elastomer was placed on the brain. An optical window (3 mm round cover glass #0, Warner Instruments) was then placed on the silicone and secured to the surrounding bone using dental cement (C&B Metabond Quick Adhesive, S380 Parkell). Another thin layer of the elastomer was placed on top of the optical window, and a GRIN lens was then secured over the optical window using dental cement attachment to the bone.

In both hippocampus and DL surgeries, a microendoscope camera (nVista, Inscopix) was used to position a magnetic baseplate. The position was chosen such that the camera was centered on the lens and focused just below the brain surface. The baseplate was secured to the bone using light-cured dental cement and then painted black (Black onyx, OPI Nail Lacquer).

### Behavioral experiments

#### Behavioral arena

Random foraging experiments were conducted in an enclosed square arena (91 × 91 × 127 cm). All surfaces of the arena were black. One wall of the arena opened to serve as a door, and another wall had a single white cue card positioned at its center. The floor was made of rubber (5812T33, McMaster-Carr). Four feeders were mounted at the center of each quadrant of the arena and dropped small pieces of sunflower seeds at irregular intervals (2.5 mg every 2-10 s). Because of the bounce of the floor, the four feeders produced a nearly uniform distribution of seeds on the arena surface.

Four infrared cameras for behavioral tracking (400300W, Qualysis) and one color camera for video monitoring (400200VC, Qualysis) were secured roughly in the corners and the center of the ceiling, respectively. During all sessions, white noise was played over a speaker to mask inadvertent room noises. Sessions lasted ∼1.5 hours, and each bird was run once per day.

#### Behavioral habituation

All behavioral experiments were performed on titmice. Prior to surgery, birds were habituated to food deprivation and handling similarly to previous work.^95^ They were first given a 4-day period of acclimation and habituation to daily handling. After this period, they were gradually habituated to food deprivation by not being given food for some amount of time at the beginning of the lights-on period each day. This food deprivation period was gradually increased from 2 h to 4 h over the course of 2 weeks. Birds were weighed daily at the end of the food deprivation period, and the length of this period was increased only if the weight was stable (fluctuations less than 0.3 g) for 3 days.

Surgery was performed on animals that had habituated to food deprivation. Following surgery animals received ad-libitum food for at least one week. They were then gradually re-habituated to food deprivation. Once birds were again maintaining a stable weight, they were introduced to the arena (see below) for daily behavioral sessions. Each session began at the end of the food deprivation period and lasted 1.5 h. After the session, birds were given ad-libitum food in their home cages for the rest of the day. This period (typically 3 days) continued until coverage of the arena was consistently greater than 98%, measured as the fraction of “pixels” covered by the animal in 1.5 h, if the arena is divided into a 40×40 grid of pixels.

Once birds were exhibiting sufficient coverage of the arena, they were habituated to the weight of the microscope and the microscope cable. Birds were first fit with a leg-loop harness,^96^ that remained permanently on the bird and whose purpose was to minimize the forces applied by the cable to the bird’s head. The harness consisted of a 3D-printed plastic attachment for the cable (21 × 9.5 × 6.5 mm) pressed against the bird’s back and two loops of elastic string (4 cm for each leg, Outus Elastic Cord, B06XJNNGMC, Amazon) that were hooked onto the bird’s thighs. Birds were first run in the behavioral arena with only the harness for 1-2 days to ensure they maintained coverage of the arena. For the remainder of the sessions, the Inscopix microscope was snapped into the magnetic headplate, and the cable at a point ∼12 cm from the microscope was attached to the harness. To further minimize the forces exerted by the cable, strands of a thin elastic string (1 mm Flat Electric Crystal Stretch String, JY001162W, Amazon) were tied to the cable to provide a low spring-constant force opposing gravity. Birds were recorded on random foraging sessions until they had completed 3-4 sessions with 98% coverage of the arena. In two early pilot birds, more sessions were acquired.

### Histology and histological imaging

Birds used for hippocampal recordings had an imprint of the GRIN lens on the surface of the brain, which could be used to verify recording coordinates. These birds were perfused after the completion of all the behavioral sessions. In birds used for DL recordings, we used an additional procedure to mark the recording location. Prior to perfusion birds were anesthetized with 1-2% isoflurane and placed into the stereotaxic apparatus. Coordinates of the lens relative to fiducial points on the skulls were obtained, and the entire implant was then removed. A pipette was lowered 0.5 mm into the brain at the center of where the lens had been placed, and 15 μL of DiI solution (5% wt/vol in DMSO, D3911, Invitrogen) was injected to mark the location. The animal was then immediately perfused.

For all perfusions, animals were administered ketamine and xylazine (10 mg/kg and 4 mg/kg respectively), then transcardially perfused with 1X PBS followed by 4% paraformaldehyde in PBS. The brains were extracted and stored in 4% paraformaldehyde in PBS for 2 days. They were then sectioned coronally into 100 μm-thick slices. Slices were stained with fluorescent DAPI (300 nM in 1X PBS, D1306, Invitrogen) and mounted in Vectashield mounting medium (H-1400-10, Vector Laboratories). Slices were imaged using a AZ100 Multizoom Slide Scanner (Nikon) using filters for DAPI and all injected fluorophores. For the images in Figure 1C, additional images were taken using a laser scanning A1R confocal Microscope (Nikon).

### Quantification and statistical analysis

All analysis was performed using custom code in MATLAB, except where specified otherwise. All code will be available for download following publication.

#### Quantification of labeled cells in anatomical tracing

To detect retrogradely labeled cells in histological slices, we used a standard Difference-of-Gaussians algorithm for blob detection. We used a series of five sigma values, logarithmically spaced from 2.5 to 12.5 μm – corresponding to blob sizes of 5 to 25 μm. The slice image was smoothed with Gaussians of each of these sigma values. Successive pairs of smoothed images were subtracted from one another to compute four difference-of-Gaussians, and these differences were stacked into a 3-dimensional matrix. Local maxima exceeding 0.02 in this matrix were detected as the centers of the blobs. If two detected blobs were overlapping, the one with the lower value was eliminated.

Slices were then annotated manually to assign detected cells to regions of interest (DL/CDL, HD, and NL). The boundaries of these brain regions were determined by comparing the DAPI-stained images to published atlases from other species.^29,31,33^ Cells were then aligned to stereotaxic coordinates by aligning slices to the reference chickadee brain (described above). To quantify the volume of a particular labeled brain region, detected cells were binned into cubic voxels with 0.1 mm side length, and the resulting volume was smoothed with the MATLAB function *imgaussfilt3* (sigma 0.05 mm). Continuous voxels containing more than 5 cells each were considered part of the labeled region.

#### Construction of 2D unfolded maps

Unfolded maps of the chickadee brain were constructed using coronal slices from one of the chickadees that had injections of the retrograde tracer in three fluorophore wavelengths at different anterior-posterior positions in the hippocampus. We adapted a standard method for creating unfolded cortical maps.^40,52^ For each slice, we marked the “medial ridge” – the dorsal-most point on the midline of the brain at which the right and left hemispheres of the forebrain contacted one another. At posterior locations (where cerebellum was present), the two forebrain hemispheres no longer contacted each other dorsally; however, a dorsal ridge continuous with the marked anterior locations was still evident and could be marked. In addition to the medial ridge, we marked the extent of retrogradely labeled cells in DL/CDL, using their approximate projections onto the brain surface. If multiple fluorophore wavelengths were present in the slice, we also marked approximate boundaries between groups of cells labeled by the different wavelengths. Finally, we marked (if present in the slice) the approximate boundary of DL and CDL on the brain surface.

All marked points in a slice were connected by tracing the outline of the brain surface. This outline was then unfolded to create a line segment with all of the marked points located at different positions on the segment. Segments obtained from different coronal slices were aligned using the medial ridge to create the 2D map. The boundaries of DL/CDL with other brain regions were determined using a published atlas of the chicken brain.^29^

To construct unfolded maps of the rat brain, we used horizontal sections from a published atlas.^54^ The surface of the entorhinal cortex was traced using the digital version of the atlas. Boundaries of MEC and LEC were also marked using this atlas. We defined the medial ridge as a “kink” evident in horizontal slices, at which the primarily-posterior wall of the cortex made a sharp turn in the anterior direction and became a primarily-medial wall. Outlines of the horizontal slices were unfolded and aligned using the medial ridge, as in chickadees. The topography of projections to the hippocampus was deduced from a published study.^40^

#### Behavioral analysis

To track the bird’s position in the behavioral arena, a spherical marker covered in infrared-reflective tape (3M Scotchlite 7610 Reflective tape, MKR-TSPE-2, B&L Engineering) was connected to the top of the Inscopix microscope. This marker was tracked using four infrared cameras recording at 300 fps (see above). The 2D position of this marker projected onto the floor of the arena and was used as the position of the animal for all analyses. Location was downsampled to 20 fps to align with the rate of image acquisition for calcium data.

To determine head direction we trained a deep neural network^97,98^ to identify the tip of the bird’s beak and the base of its neck, and drew a vector from the neck to the beak. Head direction was considered to be the angle of this vector’s projection onto the floor. The direction of the wall with the door to the arena was assigned to be the zero-degree head direction.

The animal’s speed was determined by calculating instantaneous speed and smoothing with a 1 s square window.

#### Calcium imaging

Imaging data were collected at 20 fps. Neuronal traces were extracted from raw fluorescence movies using a constrained non-negative matrix factorization algorithm intended for 1-photon calcium imaging data (CNMF_E).^61,94^ We used a multi-scale approach^99^ to extract stable fluorescent traces from long videos (∼1.5 h in our case). Before applying CNMF_E to the raw videos, we applied a motion correction algorithm.^100^ The vast majority of data contained no motion above a 1 pixel RMS shift, and any sessions that exceeded 2 pixels RMS of shift were eliminated from the analysis.

The multi-scale CNMF_E approach was run in three steps. First, data were temporally downsampled by a factor of 20. Cell footprints were found in the downsampled movie using the standard CNMF_E algorithm. These footprints were then used to extract temporal traces on segments of the non-downsampled data. Finally, the raw traces were deconvolved to detect the time and amplitude of each calcium event. To eliminate some infrequent imaging artifacts, any calcium events with an amplitude greater than 1.5 times larger than the 99th percentile of all calcium events for that cell were eliminated from all analyses. For all firing rate calculations in the paper, events were weighed by their amplitude.

#### Analysis of spatial activity

For all spatial analyses, periods where the bird was still (speed less than 5 cm/s for more than 5 s) were eliminated. Spatial firing maps were constructed by first dividing the 91×91 cm environment into 40×40 bins. For each neuron, the number of calcium events and the animal’s occupancy in each bin were calculated. The resulting two matrices were smoothed using a 13×13 Hamming window. Firing rate maps were constructed by dividing the smoothed firing rate map by the smoother occupancy and scaled to convert the units of each bin into events/s. Any bin with less than 0.1 s of occupancy after smoothing was replaced by NaN.

To detect spatially significant cells, two metrics from previous work were used: spatial information and spatial stability.^17,101^ Spatial information *I* was defined using the equation:

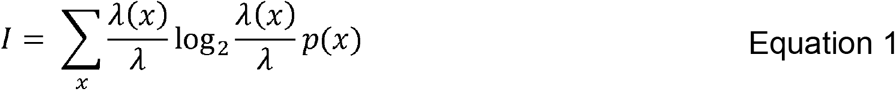

Where *x* is the index of the spatial bin, λ(*x*) is the average neural firing in bin *x*, λ is the average firing rate of the neuron, and *p*(*x*) is the probability of the animal being in the bin *x* during the session.

To compute spatial stability, we divided the session into non-overlapping 5-min segments. These segments were randomly assigned to one of two groups, such that both groups contained the same number of segments. For each group, the spatial firing map was computed, and then the correlation between the maps of each group was determined. This procedure was repeated 10 times, and then the average correlation was found.

To determine the significance of spatial information and spatial stability, we shuffled the data by circularly shifting the calcium event data by *t* timepoints relative to the behavior. For each shuffle, a random value of 0 < *t* < *T*-1 was chosen, where *T* is the total number of behavioral timepoints. The same procedures as above were used to compute spatial information and spatial stability of the shuffled data, and the shuffling procedure was repeated 500 times. Values were considered significantly high if they exceeded 99% of the respective shuffled distributions. A cell was considered “spatial” if both its spatial information and its spatial stability were significant. For calculating the normalized spatial information (Figure 8), the mean and standard deviation of the shuffled distribution were used to compute a z-score.

#### Features of spatial cells

To detect the firing fields of a cell, its firing rate map was normalized to values between 0 and 1, using the 5^th^ and 95^th^ percentiles of all the values in the rate map. A field was defined as a continuous region larger than 200 cm^2^, in which all normalized values exceeded 0.5. This procedure was used to determine the number of fields, as well as their sizes.

To determine border selectivity, we adopted a published border score.^60^ In this procedure, *c*_*M*_ was defined as the maximum fraction of any wall contacted by a single field. It was 0 if no fields touched any walls and 1 at least one of the walls was fully covered by a field. A mean firing distance *d*_*m*_ was computed by averaging, across all pixels that belonged to any field, the distances to the nearest wall, weighed by the firing rates. This value was 0 if all firing was at the exact center of the environment and 1 if all firing was on the edges of the environment. The border score *b* was then defined as:

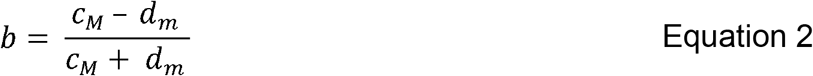

To determine if a border score was significant, shuffled values were computed as in the “Analysis of spatial activity” above. A neuron was classified as a border cell if it was significantly spatial, and its border score exceeded that of 99% of the shuffles.

To examine the optimal temporal shift of neurons, behavior was shifted between -2 s and +2 s relative to neural data in 50 ms increments. For each temporal shift, spatial information was recalculated as above. We determined the temporal shift at which the spatial information was maximal.

To attempt detecting grid cells, we measured the gridness score^62^ of each cell. First, we computed the unbiased autocorrelation of the cell’s rate map.^10^ We then defined the radius of the central peak as the distance closest to the center at which autocorrelation was either negative or exhibited a local minimum. We then considered a set of annulus-shaped samples of the autocorrelation map, where the inner radius of each annulus was the radius of the central peak, and the outer radius was varied in steps of 1 bin from a minimum of 4 bins more than the radius of the central peak to 4 bins less than the width of the environment. We then calculated the Pearson correlation of each annulus with its rotated versions at 60° and 120° (group 1) and then 30°, 90°, and 150° (group 2). The difference between the minimum of the group-1 correlation values and the maximum of the group-2 correlation values was computed, and the gridness score was defined as the maximum difference across all annulus samples. Cells whose gridness score exceeded 99% of the reshuffled samples (“Analysis of spatial activity” above) were defined as grid cells.

#### Analysis of head direction tuning and speed tuning

To detect head direction cells, we used a method based on those previously published^11^. We first divided the animal’s head direction into 9° bins. For each of these 40 bins, we defined a vector whose amplitude was the average firing rate in the bin, and whose direction was the midpoint of the bin. As in previous studies, we computed the mean vector length (MVL) as the length of the vector average of these 40 vectors. We then shuffled the data as in the “Analysis of spatial activity” above. A cell was classified as a head direction cell if its MVL exceeded 99% of the values computed on the shuffled data.

To detect speed cells, we also adopted a previously used metric.^12^ The speed score was defined as the Pearson correlation coefficient between the animal’s speed and the firing rate computed in 50-ms bins. A neuron’s speed score was considered significant if it was either below the 0.5^th^ percentile or above 99.5^th^ percentile of values from shuffled data, computed as above. This procedure detected both positively and negatively tuned cells. For displaying in figures, speed tuning curves were binned between 0 and 20 cm/s in 0.5 cm/s bins and smoothed with a 2.5 cm/s window.

#### Analysis of dash modulation

A dash onset was defined as a positive speed threshold crossings of 15 cm/s that was not preceded by another positive threshold crossing in less than 1 s. We measured peristimulus time histograms (PSTHs) aligned to dash onsets, with the firing rate smoothed by a 250-ms rolling window. Neurons modulated by dashes tended to have a peak in firing rate between 0.5 s and 1.5 s after the dash onset. We therefore quantified the strength and timing of dash modulation by detecting the maximum and the minimum of the PSTH between 0 and 2 seconds. We defined a dash score as:

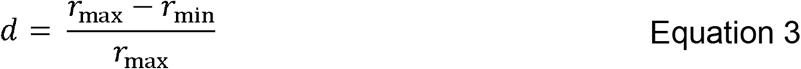

Where *r*_max_ and *r*_min_ are the firing rates at the maximum and the minimum, respectively. To determine the significance of this peak, we constructed shuffled data as in the “Analysis of spatial activity” above. Shuffling was repeated 500 times, and a cell was considered dash-modulated if the peak of its PSTH exceeded 99% of the peaks from shuffled data.

To determine whether peaks in PSTHs formed a sequence (Figure 7I), we asked whether the timing of these peaks was consistent across dashes. For each dash-modulated neuron, we randomly sorted all dashes into two equal-sized groups. We computed PSTHs separately for each group (PSTH 1 and 2), and detected the peaks of both PSTHs. For each neuron, we then computed the absolute difference in the peak timing of PSTH 1 and PSTH 2, and found the median absolute differences across neurons. We asked whether this median difference was smaller when PSTH 1 was paired with PSTH 2 from a randomly selected neuron instead of PSTH 2 from the same neuron. This procedure was repeated 1000 times, and the p value was determined as a fraction of these comparisons in which the within-neuron difference was greater than the across-neuron difference.

#### Model of multiplexed representations

Our model was largely adapted from an existing linear-nonlinear-Poisson model of entorhinal activity.^45^ To begin, the animal’s behavioral state was evaluated in 50-ms bins, synchronized with the 20 Hz acquisition of imaging data. In each bin, the animal’s position, head direction, speed, and time since last dash were averaged. These variables were discretized into *p* positional bins, *h* head direction bins, *s* speed bins, and *d* dash-time bins. As in prior analyses, any time bins where the animal’s speed was less than 5 cm/s for more than 5 s were eliminated from the model fit.

The animal’s position in the 91×91 cm environment was divided evenly into 5×5 bins (*p =* 25). For head direction, the 360° range of angles was divided into 18 identical bins (*h =* 18). The bird’s speed was discretized into 10 identical bins between 0 cm/s and 20 cm/s (*s* = 10). Time points that exceeded 20 cm/s were assigned to the last bin. Time since the most recent dash was assigned to 30 bins with logarithmically spaced edges between 0 and 35 s (*d* = 30). Any time points that were more than 35 s since the last dash were assigned to the last bin.

The animal’s state was represented by 4 sparse matrices *P, H, S*, and *D*, where each matrix had *p, h, s*, and *d* columns respectively and *T* rows, where *T* is the number of time bins in the session. For each row, these matrices had a value of 0 in all columns except the column corresponding to the value of the behavioral variable in that time bin.

Sixteen models were considered, each including some subset of the four behavioral variables (place, head direction, speed, dash modulation). The models are listed in Table S2. For each model, the matrices corresponding to the included variables were horizontally concatenated to form a matrix *X*.

Each column of *X* corresponded to a particular bin of a particular behavioral variable. We wanted to estimate the contribution of that bin to a neuron’s firing. Therefore, we defined a vertical vector of parameters *W*, whose number of rows was equal to the number of columns in *X*.

For a particular vector *W*, the estimated firing rate 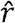 of the neuron was:

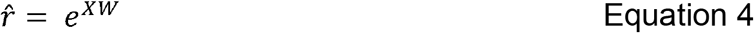

In the original model, a Poisson distribution was used to compute, in each time bin, the probability of observing a certain number of spikes. In our calcium imaging data, the number of calcium events was not always an integer, since events were weighed by their amplitudes. We therefore used a Gamma distribution Γ(*k*) instead of the Poisson distribution. Here, *k* is the mean of the distribution and the scale parameter θ is set to 1 such that the variance of the distribution (*k*θ^2^) is equal to the mean (*k*θ).

We defined *x*_*t*_ as the number of calcium events observed in time bin *t*. From the Gamma distribution, the likelihood of observing this number is given by

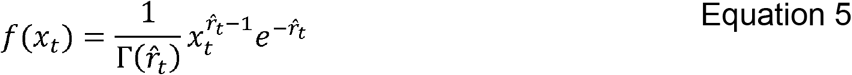

We then sought to maximize the log-likelihood across all time bins considered in the analysis. Taking the logarithm of Equation 5 and summing across all time bins, we get

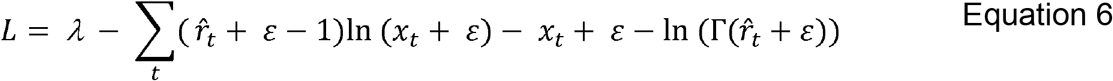

Here, ε=10^−4^ is a small value to prevent taking logarithms of 0, and λ is a smoothness parameter defined as:

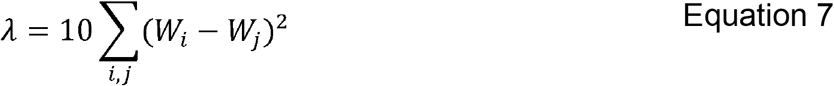

For all (*i, j*) corresponding to adjacent bins (e.g. spatial bins that share a physical border in the environment or head direction bins that are adjacent on the circle). We then computed the optimal set of parameters *W* by minimizing *L* using MATLAB function *fminunc*.

#### Model comparisons

To fit a model to a particular neuron’s activity, the behavioral session was divided into 5 time windows, which were identical in length after some of the time bins were excluded by speed thresholding. Each window was further subdivided into 10 identical sub-windows. All of the 50-ms time bins were then sorted into 10 groups, where the *n*^th^ group included all bins that were within the *n*^th^ sub-window of each of the 5 windows. For each of the 10 groups, we left that group out of the data, and fit the model to the remaining 9 groups to estimate parameters *W*. We then then computed the likelihood (Equation 6) of observing data in the left-out group given these parameters. This procedure provided us with 10 likelihood values, corresponding to each of the left-out groups. To compare any two models, we computed the average difference between these values for the two models.

To select the most appropriate model for each neuron, we first fit all models (Model #0-Model #15) to the data using the above procedure. We determined which of the one-feature models #1-#4 was the best performing model. We then asked if that model better that Model #0, which fits a single parameter to all the firing rate bins. If it was, we then found the best-performing two-feature model (Models #5-10) and compared it to the best one-feature model. We repeated this procedure for the three-feature models (Model #11-14) and the four-feature model (Model #15).

**Table S2:** Behavioral variables included by each model. *All sixteen models included in the papers. For each model, the table lists the matrices used for the subset of behavioral variables considered. These matrices are horizontally concatenated to form matrix X*.

| Model # | Components of $X$ |
| --- | --- |
| 0 | n/a (one parameter fit for all data) |
| 1 | $P$ |
| 2 | $H$ |
| 3 | $S$ |
| 4 | $D$ |
| 5 | $P, H$ |
| 6 | $P, S$ |
| 7 | $P, D$ |
| 8 | $H, S$ |
| 9 | $H, D$ |
| 10 | $S, D$ |
| 11 | $P, H, S$ |
| 12 | $P, H, D$ |
| 13 | $P, S, D$ |
| 14 | $H, S, D$ |
| 15 | $P, H, S, D$ |

