## Supplemental Figures for "An entorhinal-like region in food-caching birds"

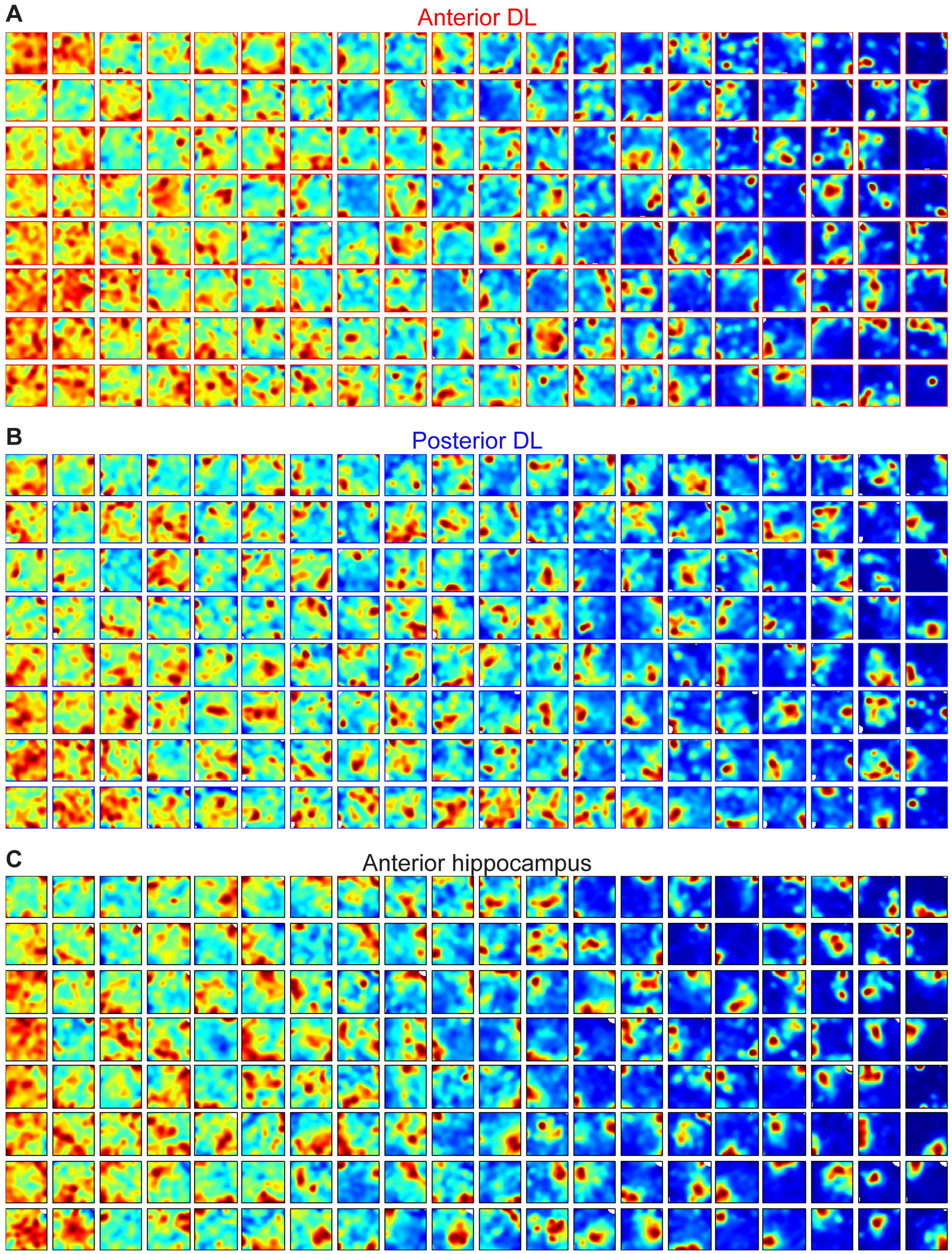

**Figure S1. Unbiased samples of spatial activity in the recorded regions**

(A) Firing rate maps from a sample of neurons recorded in anterior DL. An equal-sized group of 40 neurons was selectedfrom each of the four birds. All 160 neurons were sorted by the amount of spatial information and binned into columns, fromthe least spatial (leftmost column) to the most spatial (rightmost column). Within each column, cells were sorted top-to-bottom by their maximum firing rate. For each cell, color map ranges from 0 (blue) to 99th (red) percentile of the pixels.

(B) Same as (A), but for posterior DL.

(C) Same as (A), but for anterior hippocampus.

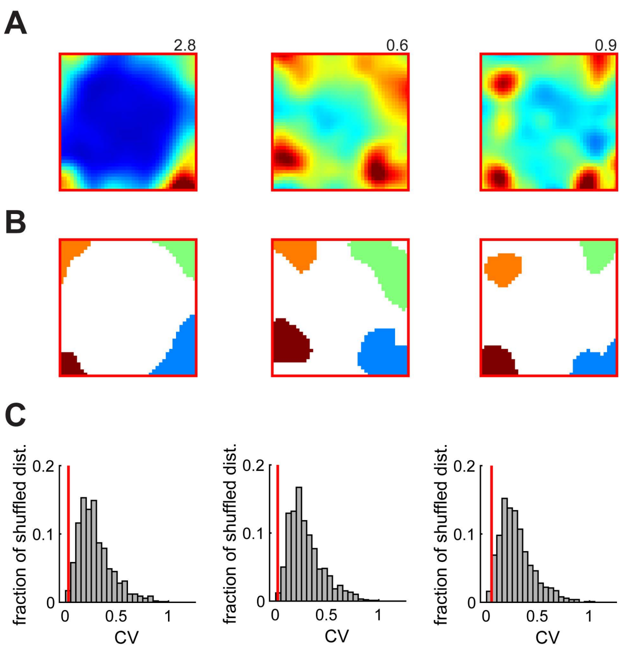


**Figure S2. Ordered arrangement of fields in anterior DL**

(A) Firing rate maps examples of three cells in anterior DL, each showing four firing fields.

(B) Firing fields detected in the rate maps shown in (A).

(C) Analysis of spacing between the fields for each of the cells in (A). The position of each field was considered to be its center of mass. For each field, the two nearest neighbor fields we found, and the average distance to those two neighbors was calculated. This process produced a single distance value for each field. The coefficient of variation (CV) of these values was then computed for the cell. We asked whether this value was smaller than expected by chance. To answer this, we generated random field locations from a uniform distribution spanning the entire environment, not allowing the distance between any pair of locations to be smaller than 25 cm (the smallest distance found in the actual data). The process was repeated 1000 times, and the CV of distances was computed each time, as for the actual data. A cell was considered to have a significantly regular arrangement of fields if its CV was smaller than that of 95% of the random field arrangements. In the figure, histogram shows the distribution of CVs for randomly-generated data; vertical line shows the value for the actual data. All three examples are significant. Across the population in anterior DL, there were 36 cells with four or more fields; 9 of these cells (25%) were significant. Though based on published analysis in flying bats,[23] our analysis uses different thresholds and inclusion criteria. The reason is that our recordings were in a much smaller environment and in 2D, rather than 3D; our cells generally had fewer fields than those recorded in the MEC of flying bats.

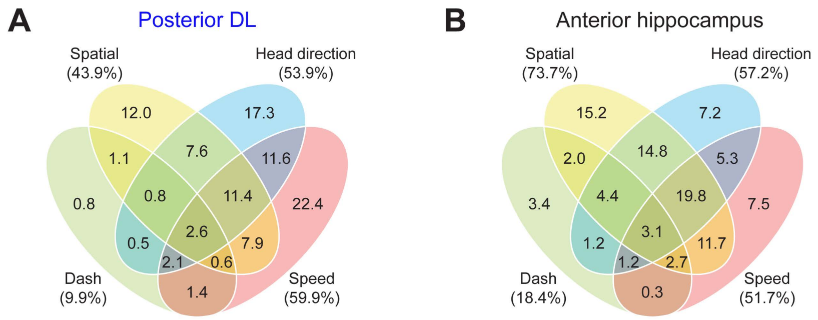


**Figure S3. Mixed representations in posterior DL and anterior hippocampus**

(A) Venn diagram of posterior DL cells, classified according to what subset of the four variables they were modulated by. The diagram classifies 251 cells that were considered by the model to be modulated by at least one variable; an additional 23 cells were not modulated by any variable.

(B) Diagram as in (A), but for anterior hippocampus. The diagram classifies 425 cells; an additional 38 cells were not modulated by any variable.
